# Accelerating Antibody Development: Sequence and Structure-Based Models for Predicting Developability Properties through Size Exclusion Chromatography

**DOI:** 10.1101/2025.02.02.636157

**Authors:** A N M Nafiz Abeer, Mehdi Boroumand, Isabelle Sermadiras, Jenna G Caldwell, Valentin Stanev, Neil Mody, Gilad Kaplan, James Savery, Rebecca Croasdale-Wood, Maryam Pouryahya

**Affiliations:** Data Science and Modelling, BioPharmaceuticals R&D, AstraZeneca; Department of Electrical and Computer Engineering, Texas A&M University; Biologics Engineering, Oncology R&D, AstraZeneca; Dosage Form Design and Development, BioPharmaceuticals R&D, AstraZeneca

## Abstract

Experimental screening for biopharmaceutical developability properties typically relies on resource-intensive, and time-consuming assays such as size exclusion chromatography (SEC). This study highlights the potential of in silico models to accelerate the screening process by exploring sequence and structure-based machine learning techniques. Specifically, we compared surrogate models based on pre-computed features extracted from sequence and predicted structure with sequence-based approaches using protein language models (PLMs) like ESM-2. In addition to different end-to-end fine-tuning strategies for PLM, we have also investigated the integration of the structural information of the antibodies into the prediction pipeline through graph neural networks (GNN). We applied these different methods for predicting protein aggregation propensity using a dataset of approximately 1200 Immunoglobulin G (IgG1) molecules. Through this empirical evaluation, our study identifies the most effective in silico approach for predicting developability properties for SEC assays, thereby adding insights to existing screening efforts for accelerating the antibody development process.

## 1. Introduction

Monoclonal antibodies (mAbs) are effective therapeutic proteins due to their high specificity, versatility, and efficacy in targeting a wide range of diseases. They enable targeted therapy with minimal off-target effects, applicable in oncology, autoimmune, infectious, cardiovascular, and metabolic diseases (Kaplon & Reichert, 2018). Advances in antibody engineering, such as humanization and affinity maturation, have enhanced their clinical efficacy and reduced immunogenicity, making mAbs safer and therapeutically effective for long-term use (Saxena et al., 2009). With over 145 antibody-based drugs approved by the FDA and many more in clinical trials, their impact on modern medicine is substantial and continues to grow (Strohl, 2024; Ecker et al., 2015).

The developability properties of antibodies are crucial for their transition from early-stage discovery to large-scale manufacturing. Since the antibody design process prioritizes the binding activity in neutralizing the target protein, the resulting antibodies may end up with unfavorable biophysical attributes hindering its progression into the next development stages (Jain et al., 2017; 2023). Hence, it is extremely beneficial to screen antibodies not only for target specificity and binding affinity but also for physicochemical and biopharmaceutical properties such as solubility, stability, aggregation propensity, and manufacturability (Venkatesh & Lipper, 2000). These properties significantly influence the antibody’s success in later development stages, including clinical trials and commercialization (Kola & Landis, 2004). Poor developability can lead to high attrition rates, increased costs, and extended timelines due to formulation challenges, instability, or immunogenicity (Sun et al., 2004). Systematic early evaluation of these properties can streamline development, advancing only the most promising candidates (Serajuddin, 2007). This proactive approach also aids in designing robust manufacturing processes to produce high-quality therapeutic antibodies at scale, reducing late-stage failures and ensuring a more efficient path to commercialization (Garad, Sudhakar, 2004; Saxena et al., 2009).

Various assays and methods, such as size exclusion chromatography (SEC) (Mori & Barth, 1999), dynamic light scattering (DLS) (Stetefeld et al., 2016), differential scanning calorimetry (DSC) (Johnson, 2013), and isoelectric focusing (IEF) (Righetti, 1983), are employed to measure the physicochemical attributes of antibodies, serving as proxy evaluations for their developability characteristics. These measurements are utilized to detect aggregates, assess conformational stability, evaluate chemical degradation, and determine both the isoelectric point and charge isoform content, all of which impact the molecule’s efficacy, solubility, and overall chemical stability. (Saxena et al., 2009; Venkatesh & Lipper, 2000; Jain et al., 2017). In our work, we have considered the SEC assay which offers unique advantages in the purification and characterization of biomolecules. This chromatographic method, which separates analytes based on molecular size, plays a critical role in ensuring the purity, efficacy, and safety of therapeutic proteins. SEC allows for the effective separation of desired therapeutic proteins from aggregates and impurities that may affect the drug’s safety and efficacy. Furthermore, SEC is crucial for the detailed characterization of protein biotherapeutics, providing insights into their molecular weight distribution, aggregation state, and stability, which are essential parameters for regulatory approval and clinical success (Fekete et al., 2014; D’Atri et al., 2024). The technique’s ability to operate under mild conditions without altering the biological activity of the molecules makes it particularly valuable for the analysis of sensitive biologics (Hong et al., 2012), (Chakrabarti, 2018).

While the experimental assays like SEC provide valuable insights into the development of biologics, they are often time-consuming and costly. The high-throughput screening methods required to evaluate a large number of candidates can be resource-intensive, requiring significant investment in both equipment and materials (Balbach & Korn, 2004). Machine learning and in silico approaches can significantly enhance the prediction of antibody developability by leveraging medium to high-throughput datasets. These methods aid in early-stage drug development, enabling the rapid selection of lead candidates and potentially reducing the need for extensive high-throughput analytical measurements. However, the limited amount of experimental assay datapoints poses a critical challenge in building a reliable surrogate model for predicting the developability property of interest. The existing efforts (Bailly et al., 2020; Waight et al., 2023; Rai et al., 2023; Park & Izadi, 2024; Rollins et al., 2024a) primarily involve processing the sequences (with or without 3D protein structure) to compute the protein descriptors that are utilized by machine learning models for prediction. In addition to the computational burden of generating the features from structures, the performance of this approach is sensitive to the feature selection process. The protein language models (PLMs) offer a faster and more efficient alternative to utilize the structure information of the proteins. Leveraging the large pool of protein sequences, the PLMs learn to incorporate the structural information implicitly into the sequence embedding (Rao et al., 2021). With the advancement of protein language models, there have been efforts (Villegas-Morcillo et al., 2021; Wang et al., 2022) to predict the protein property from the sequence embedding learned by the pre-trained PLM. Since the PLMs work on the sequence representations of the antibodies, this approach has the potential for an accurate and faster screening pipeline by removing the need for prediction of structure and processing of structural features. On the other hand, several works (Wang et al., 2022; Widatalla et al., 2023; Rollins et al., 2024b) explicitly leverage the 3D protein structures along with the protein language model for the prediction of protein properties. In the absence of experimental structure data, this approach relies on either the protein folding tools like AlphaFold2 (Jumper et al., 2021), or homology modeling to build the structure from the protein sequence.

The performance of the above-mentioned approaches for predicting the protein properties varies across different types of assays. In this work, we have investigated the application of these different prediction approaches for the SEC assay. Specifically, we have considered the classification task of two developability properties – monomer content and difference in retention time of the IgG1 molecules to a reference sample. To summarize our work:

- We performed experiments with four prediction pipelines – one based on sequence and structure-based features, three others leveraging the protein language model and graph neural network – to select the best performing configuration (e.g. which features/PLM/GNN to use etc.) under each pipeline.
- We have identified the best prediction strategy for two SEC properties of interest based on the hold-out test set performance by these four pipelines.
- Furthermore, we have assessed the impact of two different protein structure prediction tools on the developability prediction pipeline.

## 2. Methodology

### 2.1 Problem statement

Experimental assays for screening antibodies, such as SEC, can characterize multiple attributes of the molecule under consideration (Section 3.1). For one of such properties, we assume to have a dataset 𝒟 = {***s***^(*i*)^,*o*^(*i*)^} where ***s*** denotes the sequence representation of the antibody, i.e. both heavy and light chains. The attribute *o* ∈ ℝ corresponds to the observation value directly obtained through the assay. During the screening stage, the sample is classified as desirable or problematic by comparison with a pre-defined specification, usually based on developability requirements. By denoting this process as f_screen_ we can get the binary label, *y* ∈ {0, 1}, i.e. desirable/problematic for each sample as:

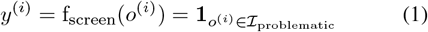

**1** is an indicator function which labels the sample 1 (problematic) when the corresponding observation falls within the interval _problematic_. This interval expectedly varies across different properties (Figure 1).

**Figure 1:**
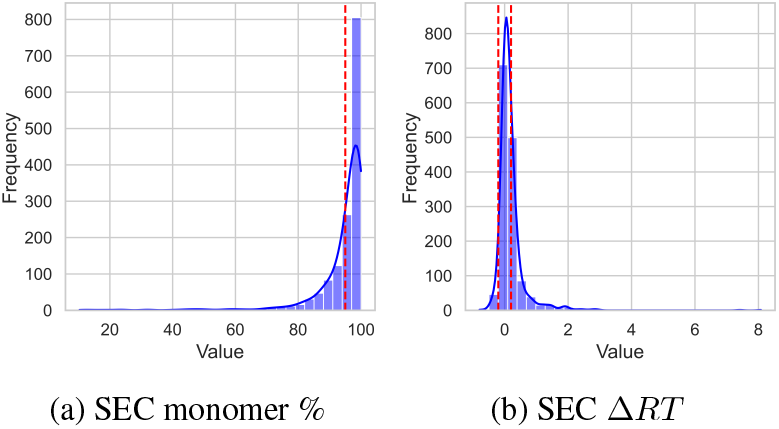
Data distribution of SEC monomer % and ΔRT. Problematic antibodies are characterized by lower monomer content and Δ*RT* values further from zero, as indicated by the red dashed lines in the figure.

Given the data with binary labels, 𝒟 _*bin*_ = {***s***^(*i*)^, *y*^(*i*)^}, our goal is to build a classifier to identify an Immunoglobulin G (IgG1) molecule as a desirable (developable) or problematic sample from information embedded in ***s***. Once we have such a classifier trained, this can serve as a surrogate for the experimental assay in the in silico screening process.

### 2.2. Prediction Pipeline

The primary data modality in our study is the sequence representation. ***s*** = (***s***_heavy_, ***s***_light_) of the antibody. All molecules in our dataset belong to the IgG1 subclass and share a similar constant region. Consequently, our predictions concentrate exclusively on the impact of sequences within the variable fragments (Fvs). Through the application of the AlphaFold2 (AF2) (Jumper et al., 2021) or similar protein folding tools, one can also predict the 3D structure folded from the sequence information alone, adding another modality to our prediction pipeline. While explicit incorporation of structure information provides richer information than the sequence for predicting protein properties, one also needs to consider possible errors propagated from the protein structure prediction tool’s inaccuracy.

In this work, we have considered the following four approaches (illustrated in Figure 2) for building the prediction network for the developability property of interest by leveraging different combinations of both modalities. Details of each pipeline are discussed in the subsequent sections.

**Figure 2:**
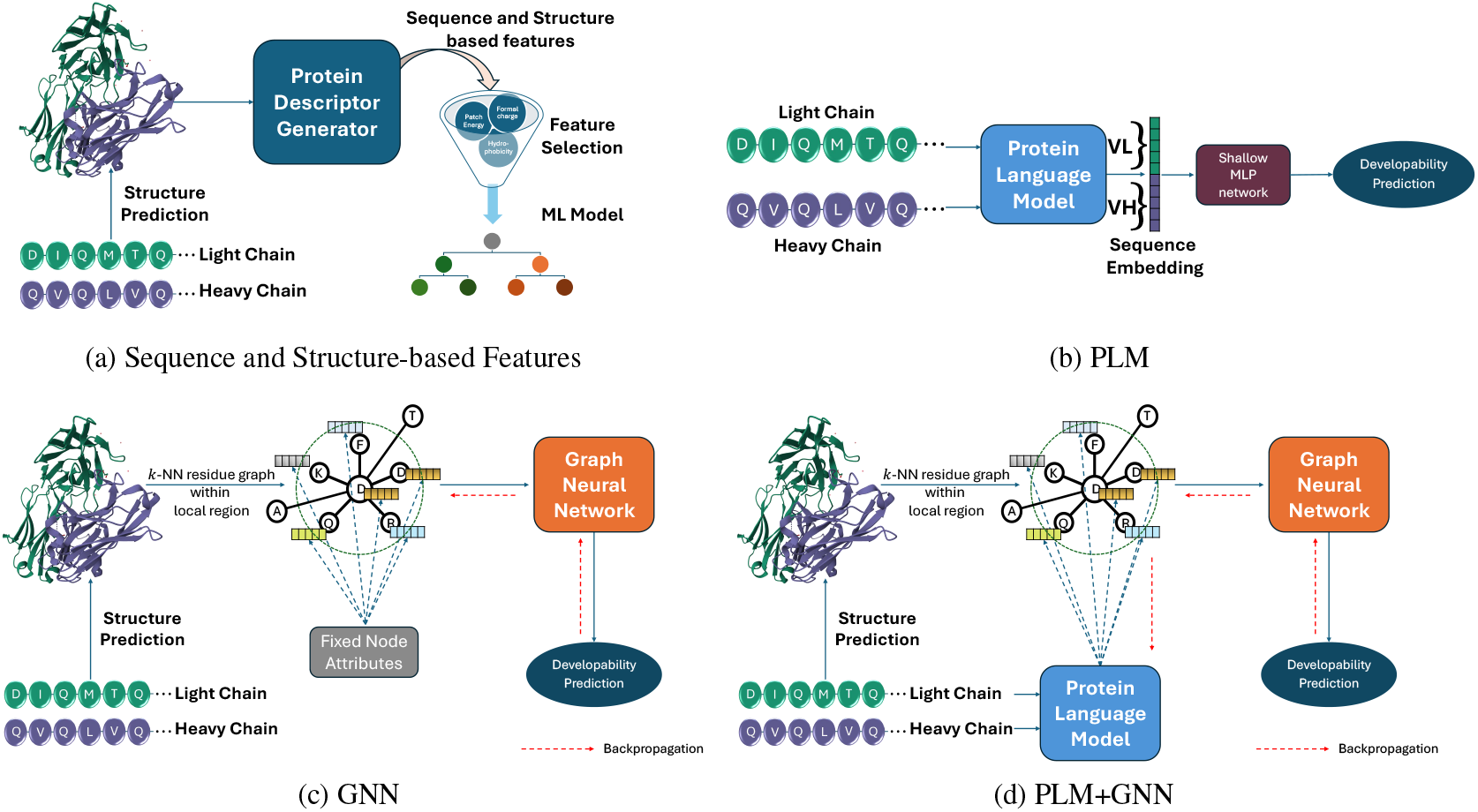
Developability property prediction workflow. All four pipelines except PLM (b) leverage the predicted 3D protein structure into their prediction network by either protein descriptor generator (a) or graph neural network (c and d). Note that GNN pipeline (c) utilizes the fixed node attributes which can include the embeddings from a pre-trained protein language model. In the PLM+GNN (d), we allow the PLM to be updated jointly with the GNN during training of the prediction network.

- **Sequence and Structure-based Features**: An ML classifier predicts the target from the selected protein features, processed by a protein descriptor generator from the antibody sequence and its predicted structure.
- **PLM Pipeline**: The pipeline utilizes only the sequence information through a combination of a protein language model (PLM) and a prediction head. The latter component is a shallow multilayer perceptron (MLP) network that predicts the property from sequence embedding projected by the PLM.
- **GNN Pipeline**: The graph neural network (GNN) predicts the property from the amino acid (AA) graph constructed from the predicted structure.
- **PLM + GNN Pipeline**: This incorporates the residue embedding from the protein language model into the GNN pipeline as the node attributes of the AA graph and the combined network is trained jointly.

#### 2.2.1. Prediction UTILIZING SEQUENCE AND STRUCTURAL FEATURES

We used Schrödinger software to extract the molecular properties from the predicted AF2 static structure (Schrödinger, LLC). Schrö dinger software offers an extensive suite of computational tools for molecular modeling, simulation, and extraction of protein descriptors. In our workflow, we utilized AF2 instead of Schrödinger’s homology modeling and employed Schrödinger solely for extracting molecular properties and biophysical features. Schrö dinger provides an exhaustive list of patch-level sequence- and structurebased protein properties, including positive and negative charges, hydrophobicity, and aggregation propensities. To ensure we selected the most informative features for our final model, we utilized a combination of unsupervised and supervised feature selection Section 3.4.1. Finally, an Extra Trees classifier (Geurts et al., 2006) was applied to the selected features for predicting the developability properties.

#### 2.2.2. PLM BASED PREDICTION NETWORK FROM SEQUENCE REPRESENTATION

The core idea of the PLM pipeline is to utilize the sequence embedding of an antibody sequence from the protein language model in combination with a simple prediction head network to classify the developability property. For the heavy chain and light chain in the antibody sequence, we consider separate instances of the protein language model, *ϕ*_PLM, H_ and *ϕ*_PLM, L_ respectively. We first compute the hidden states of each chain using corresponding instance of PLM. Specifically, we denote residue embedding matrix ***E***_H_, ***E***_L_ as the last hidden state of all tokens of each chain. Note these embedding matrices can include the hidden state of special tokens added for the PLM. Next, we compute the sequence-level representation, i.e. ***e***_H_, ***e***_L_ for each chain by pooling its embedding matrix. We have explored two aggregation techniques to do this – mean pooling where the residue embeddings (excluding any special tokens) are averaged and CLS pooling where embedding for the special token [CLS] is considered as the sequence-level embedding.

Finally we concatenate the sequence embeddings ***e***_H_ and ***e***_L_ and pass it to a shallow MLP network *ϕ*_MLP_ to predict the class label (i.e., the logits) for the sequence ***s***.

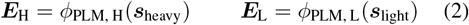

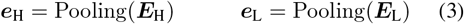

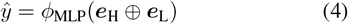

In case of general protein language model like ESM2 (Lin et al., 2023; 2022), *ϕ*_PLM, H_ and *ϕ*_PLM, L_ are initialized at the pre-trained weights. One can also consider the antibody chain specific pre-trained models like AbLang-1 (Olsen et al., 2022). When both instances are frozen at their pretrained model weights, and we learn the prediction head *ϕ*_MLP_, we denote this as “fixed PLM” approach in our work. On the other hand, we can also finetune those PLM instances while training the prediction head by allowing the gradient information backpropagated to the layers of the PLMs. We have investigated two ways of performing this finetuning – full parameter finetuning and low rank adaptation (LoRA) (Hu et al., 2022) technique. The latter approach learns lowrank transformation matrices (for the attention networks in our work) that work in conjunction with pre-trained weights to adapt the model for a specific task.

#### 2.2.3. GNN BASED PREDICTION NETWORK LEVERAGING STRUCTURE

To leverage the protein structural information explicitly in the prediction pipeline, the GNN pipeline begins with the construction of the amino acid graph for the antibody sequence ***s*** from the predicted 3D structure. For each residue in the sequence ***s***, the top *k* nearest residues are considered to be its neighbors based on the coordinates of *C*-*α* of the residues. Up to this step, the procedure is similar to the approaches of (Ingraham et al., 2019; Jing et al., 2021; Wang et al., 2022; Widatalla et al., 2023). In our work, however, we add a refinement step by pruning residues that lie outside a predefined local region. Hence, in the resulting AA graph 𝒢, the edge from node *j* to node *i* exists if residue *j* is in the *k*-nearest neighbors of residue *i* and distance of the edge, ||***e***_*j*→*i*_||_2_ is lower than *r*_thr_ Å . In our work, we set *r*_thr_ = 9 which results in around 13 neighbors on average for the nodes of AA graphs in our dataset.

The *m* layers of GNN, denoted as *ϕ*_GNN_ projects the node attribute matrix 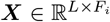 of AA graph 𝒢 for sequence ***s*** with *L* residues into the hidden embeddings matrix 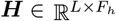. *F*_*i*_ and *F*_*h*_ are the dimension of node attributes and hidden representation respectively. The prediction head *ϕ*_MLP_ transforms the pooled embedding from ***H*** into the class probability for the antibody sample.

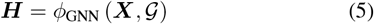

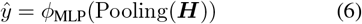

For the node attributes of the graph, we have considered the one-hot encoding, amino acid properties from (Jamasb et al., 2022; Gasteiger et al., 2003), node embedding from the pretrained variational graph autoencoder (VGAE) of (Nguyen & Hy, 2024) and residue embedding from pre-trained ESM2 (8M) model. In case of the VGAE and ESM2 (8M), the weights of these models are frozen at the pre-trained values in the entire training process. To evaluate the effectiveness of GNN in processing the AA graph, three different GNNs are considered – geometric vector perception (GVP) (Jing et al., 2021), graph attention network (GAT) (Veličković et al., 2018) and graph isomorphism network (GI) (Xu et al., 2018). The main difference between these three is that the GVP leverages the edge features in the message-passing updates while the other two do not. Also, the GVP considers the scalar and vector features for the nodes and edges where the scalar features of the nodes are based on dihedral angles and one of the four node attributes mentioned earlier. In case of other two GNNs, only the node attributes are utilized.

After 3 layers of GNN, the updated node embeddings are aggregated to get the global embedding for the entire graph. Here, we have considered the mean pooling technique which takes an average of node embeddings across all nodes of the graph. In summary, we have explored 4 choices of node attributes and 3 choices of GNN for each developability property.

#### 2.2.4. Prediction NETWORK LEVERAGING PLM AND GNN

In terms of architectural similarity, the PLM+GNN pipeline is the same as the GNN-based prediction network where the node attributes of the amino acid residue graph are assigned by the pre-trained protein language model. While in the GNN pipeline, we only learn the GNN modules during training, the PLM+GNN pipeline additionally facilitates the adaptation of the parameters of the PLM leveraging the gradients at the node attributes backpropagated from the GNN modules as in (Wang et al., 2022). For an antibody sequence ***s***, first the PLM module *ϕ* _PLM_ computes the node attribute matrix ***X***. The rest of the steps are similar to the GNN pipeline where ***X*** is transformed to the prediction through *ϕ*_GNN_ and *ϕ*_MLP_ by using corresponding amino acid graph 𝒢.

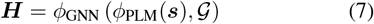

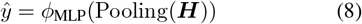

For this approach, we have considered 3 PLMs – (ESM2 (8M) (Lin et al., 2022), AbLang-1 (Olsen et al., 2022) and AbLang-2 (Olsen et al., 2024)). For the ESM2 (8M), we allow the gradient flow through its all layers excluding the embedding layer. In the case of finetuning of AbLang-1 and AbLang-2, the first 6 layers are frozen at their pre-trained weights while the rest of the layers are updated based on the gradient backpropagated from the GNN module. In addition to the global mean pooling of the GNN pipeline, we have investigated the effectiveness of universal pooling (Navarin et al., 2019; Widatalla et al., 2023) over the mean pooling technique. Although the sequences in our work are not processed using the multiple sequence alignment (MSA) technique, our exploration attempts to analyze the robustness of this pooling technique for 3 different choices of GNN.

## 3. Result and Discussion

### 3.1. High-performance Size exclusion chromatography assay

HPSEC assays help determine the purity of protein samples by measuring the percentage of protein monomer (main product), HMWFs (higher molecular weight forms) and LMWFs (lower molecular weight forms). The chromatogram results from SEC assays typically display several peaks corresponding to detected species. Peaks appearing before the expected monomer peak are classified as HMWFs, while those appearing after are identified as LMWFs. This interpretation is crucial for understanding the composition and quality of the protein sample. The chromatograms also provide the retention time of peaks, which indicate the elution time of the protein in the chromatography columns. The SEC retention time of the monomer is primarily influenced by its molecular weight, but it is also affected by other molecular properties, including charge, hydrophobicity, and self-association (Giddings, 1965), suggesting that SEC can offer a multifaceted view of the molecule’s behavior (Arakawa & Timasheff, 1985; Jain et al., 2017). To normalize retention times of monomer content across multiple studies utilized in this paper, Δ*RT* is considered, which compares the retention time of the monomer protein to a reference protein, NIP228, with a known SEC retention time of approximately 8.47 minutes. This normalization is essential for consistent analysis across different studies (Podwojski et al., 2009; Hong et al., 2012). In this work, we focus on predicting the percentage peak area for the monomer product and the delta retention time of the monomer peak. Since these assays are primarily used to distinguish problematic molecules from developable ones using well-defined thresholds, we concentrate mainly on a classification problem to differentiate between the two.

### 3.2. Dataset and train-test split

The SEC dataset, consisting of around 1200 IgG1 molecules, was collected from multiple internal studies. We considered the monomer content percentage and the Δ*RT* of the antibodies relative to a reference antibody, NIP228, as previously described. Duplicate samples, defined as multiple assay observations for an identical antibody sequence, were removed for each property. The processed dataset was then divided into training (90%) and test (10%) partitions. Due to the dataset’s collection from various studies, the sequences exhibit considerable diversity, with a median of over 100 mutations between pairs. Despite this diversity, we still specifically chose the test set to remain diverse, based on clusters formed according to sequence similarities (Levenshtein distance with representative members of clusters chosen randomly (Levenshtein, 1965; Berger et al., 2021)). This selection also ensured a similar stratified distribution of the two classes as observed in the train set. The test split serves as a hold-out set for evaluating the efficacy of the prediction pipelines (Section 3.4.2).

### 3.3. Processing of 3D protein structure

In this work, we utilized two state-of-the-art computational tools for predicting protein structures: AlphaFold 2 and ImmuneBuilder. AlphaFold 2, developed by DeepMind, has revolutionized the field by accurately predicting protein structures from amino acid sequences, applicable across a wide range of proteins (Jumper et al., 2021). ImmuneBuilder, on the other hand, is tailored for rapid and accurate predictions of immune protein structures, such as antibodies and T-cell receptors. Noted for its speed, ImmuneBuilder is significantly faster than AlphaFold 2 (Abanades et al., 2023) and more suitable for high-throughput screening of large numbers of antibodies. Therefore, we conducted a comprehensive comparison of the structure prediction models in Section 3.4.3.

### 3.4 Experiments

#### 3.4.1. Selecting BEST COMBINATION FOR EACH PIPELINE

We have explored different combinations for each of the four prediction pipelines discussed in Section 2.2. In each pipeline, we have run 10 trials of 10-fold cross-validation on the SEC dataset using different combinations and selected the best configuration based on the average prediction performance on the validation splits. Appendix A describes the pipeline-specific procedures and explains the corresponding cross-validation results in detail.

#### 3.4.2. Results ON TEST SET

From the cross-validation experiments of Section 3.4.1, we have only identified the best combination for each pipeline. Next, we have trained each best combination with 10-fold cross-validation with the training split and evaluated the trained models’ performance on the hold-out test set. Specifically, we split the training data (90% of the SEC dataset) into 10-fold cross-validation splits, and we pick the model with best performance (accuracy) on the validation split in each of 10 folds. These 10 trained models for each pipeline are applied on the hold-out test set (Section 3.2) to measure the predictive performance. Tables 1 and 2 show the average and standard deviation of performance metrics across these 10 models for each pipeline corresponding to monomer percentage and Δ*RT* respectively.

**Table 1:** Prediction performance on the hold-out test set for SEC monomer %. The majority class (label 0) predictor has an accuracy of 71%. ESM2 (8M) model fine-tuned via LoRA technique produces the sequence embedding for “PLM” pipeline. For “GNN” pipeline, the embedding from pre-trained VGAE is used with 3 layers of GVPs. For “PLM+GNN”, a combination of AbLang1 and GVP is trained in an end-to-end fashion. The best-performing pipeline is highlighted for each metric based on their average performance over hold-out test set for 10 models from 10-fold cross-validation.

| Prediction Pipeline | Accuracy | F1 | Sensitivity |
| --- | --- | --- | --- |
| PLM | 0.75 (0.03) | **0.54** (0.04) | **0.52** (0.05) |
| GNN | 0.74 (0.02) | 0.49 (0.03) | 0.44 (0.04) |
| PLM + GNN | 0.75 (0.01) | 0.50 (0.04) | 0.43 (0.06) |
| Sequence + Structure based features | **0.77** (0.01) | 0.49 (0.04) | 0.38 (0.05) |

**Table 2:** Prediction performance on the hold-out test set for SEC Δ*RT* . The majority class (label 0) predictor has an accuracy of 69%. The “PLM” pipeline utilizes the ESM2 (8M) model obtained after the full parameter fine-tuning with the training set. For “GNN” pipeline, the embedding from pre-trained ESM2 (8M) is used with 3 layers of GVPs. For “PLM+GNN”, a combination of AbLang1 and GVP is trained in an end-to-end fashion. The best-performing pipeline is highlighted for each metric based on their average performance over hold-out test set for 10 models from 10-fold cross-validation.

| Prediction Pipeline | Accuracy | F1 | Sensitivity |
| --- | --- | --- | --- |
| PLM | 0.77 (0.02) | 0.60 (0.04) | 0.56 (0.06) |
| GNN | **0.80** (0.03) | **0.64** (0.07) | **0.58** (0.10) |
| PLM + GNN | 0.78 (0.04) | 0.61 (0.13) | 0.57 (0.15) |
| Sequence + Structure based features | **0.80** (0.01) | 0.59 (0.03) | 0.46 (0.05) |

For both properties, the pipeline utilizing sequence and structure-based features achieves the highest accuracy while performing poorly in selecting problematic molecules (class label 1) as evidenced by the low value in sensitivity. The other three pipelines show slightly lower accuracy in SEC monomer percentage and out of these three, the PLM pipeline produces better F1 score and sensitivity. This result is particularly interesting for high throughput screening since this PLM pipeline predicts the developability property from the sequences, making it much faster than “sequence + structure-based features” which requires computationally time-consuming feature processing by the Schrödinger suite. In the case of the SEC Δ*RT*, the GNN pipeline and the feature-based pipeline have similar accuracy but the former shows better sensitivity. When the structural information is combined with the PLM approach resulting in PLM+GNN, the average performance is slightly increased compared to the PLM approach but this is achieved at the expense of larger variation in performance across different models learned in each of the 10 folds.

All four pipelines have a similar standard deviation in F1 score (and sensitivity) for the SEC monomer percentage property. On the other hand for SEC Δ*RT*, the GNN and PLM+GNN pipelines show comparatively larger fluctuation in similar performance metrics. For example, the standard deviations in sensitivity for these two pipelines are 0.10 and 0.15 respectively (Table 2) which are 2-3 times larger than the other two pipelines. Given that the PLM approach shows similar fluctuation as the pipeline with sequence and structure-based features, this higher variation in the performance metrics possibly originates from the sensitivity of GNN components to AA graphs seen in the 10 fold crossvalidation training. Additionally, the difference in this trend between the monomer percentage and Δ*RT* indicates that the structural information of the antibody molecules may have more importance in the predictive performance of the latter property.

#### 3.4.3. Ablation STUDY FOR DIFFERENT STRUCTURE PREDICTION TOOLS

The results in Tables 1 and 2 are for the experiments utilizing the predicted structure from AlphaFold2. Despite having a high structure prediction performance, AlphaFold2 can be a source of bottleneck in the high throughput screening process due to its longer structure prediction time. In this section, we used a faster structure prediction tool, ImmuneBuilder (Abanades et al., 2023) for the three pipelines that use antibody structure in predicting the developability properties. By comparing with the previous results (from AlphaFold2), we assessed the robustness of the developability prediction pipeline to the predicted structures.

For three developability prediction pipelines – Sequence + Structure based features, GNN, PLM+GNN – we have repeated the hold-out test set experiment from Section 3.4.2. The models under each pipeline have the same architecture as in Section 3.4.2 but are trained with the datapoints where the structures are predicted via ImmuneBuilder. Tables 3 and 4 have these hold-out set performance metrics for monomer % and Δ*RT* respectively. We have also shown the results with AlphaFold2 (from Tables 1 and 2) for reference. For both properties, we observe a decline in the sensitivity (and F1 score) for Sequence + Structure based features and GNN pipelines when we replace AlphaFold2 with ImmuneBuilder as the antibody structure prediction tools. The performance of PLM+GNN pipeline remains similar for SEC monomer % while we see a more stable performance trend for SEC Δ*RT* . With AlphaFold2, the GNN pipeline is the best performing pipeline for Δ*RT* but has a relatively large performance variation. In the PLM+GNN pipeline for Δ*RT*, the ImmuneBuilder not only works as a faster structure prediction tool but provides a comparable performance consistently (0.62(0.04) vs 0.61(0.13) with AlphaFold2).

## 4. Conclusion

Our work has explored four different prediction approaches for two developability properties from the SEC assays: monomer percentage and Δ*RT*. We aimed to provide promising in silico models that can be used to screen less developable molecules at an early stage without the need for time- and material-intensive experimantal methods. We investigated high-throughput models that require only sequences (protein language models) as well as models that require protein structure prediction and feature extraction from predicted structures, which are computationally more demanding. Specifically, we examined whether protein language models and graph neural networks can effectively utilize sequence and structural information to achieve performance comparable to methods employing explicit structural and sequence features calculated by Schrö dinger software. For both properties, the performance on the hold-out test set shows that the latter approach achieves higher accuracy but suffers from considerable degradation in its ability to identify molecules violating the developability constraint. The best-performing pipeline (in terms of F1 score) for monomer percentage is the protein language model (PLM) approach, which offers a promising opportunity for high-throughput screening due to its faster inference time from the sequence of the antibody chains. Further comparison of variations in the performance metrics for Δ*RT* and monomer percentage indicates the relative importance of structural information in predicting these two properties. Finally, we have identified (in Section 3.4.3) the potential of the PLM+GNN pipeline as a high-throughput screening tool for Δ*RT*, with ImmuneBuilder (replacing AlphaFold2) as the protein structure prediction tool.

**Table 3:** Variation in predictive performance for SEC monomer % due to different structure prediction tools. The neural network architectures for three prediction pipelines are the same as in Table 1.

| Prediction Pipeline | Structure Prediction Tool | Accuracy | F1 | Sensitivity |
| --- | --- | --- | --- | --- |
| GNN | AlphaFold2 | 0.74 (0.02) | 0.49 (0.03) | 0.44 (0.04) |
|  | ImmuneBuilder | 0.75 (0.02) | 0.46 (0.04) | 0.38 (0.04) |
| PLM + GNN | AlphaFold2 | 0.75 (0.01) | 0.50 (0.04) | 0.43 (0.06) |
|  | ImmuneBuilder | 0.74 (0.02) | 0.49 (0.05) | 0.44 (0.08) |
| Sequence + Structure based features | AlphaFold2 | 0.77 (0.01) | 0.49 (0.04) | 0.38 (0.05) |
|  | ImmuneBuilder | 0.76 (0.01) | 0.45 (0.04) | 0.34 (0.04) |

**Table 4:** Variation in predictive performance for SEC Δ*RT* due to different structure prediction tools. The neural network architectures for three prediction pipelines are the same as in Table 2.

| Prediction Pipeline | Structure Prediction Tool | Accuracy | F1 | Sensitivity |
| --- | --- | --- | --- | --- |
| GNN | AlphaFold2 | 0.80 (0.03) | 0.64 (0.07) | 0.58 (0.10) |
|  | ImmuneBuilder | 0.78 (0.03) | 0.59 (0.07) | 0.52 (0.10) |
| PLM + GNN | AlphaFold2 | 0.78 (0.04) | 0.61 (0.13) | 0.57 (0.15) |
|  | ImmuneBuilder | 0.79 (0.01) | 0.62 (0.04) | 0.55 (0.06) |
| Sequence + Structure based features | AlphaFold2 | 0.80 (0.01) | 0.59 (0.03) | 0.46 (0.05) |
|  | ImmuneBuilder | 0.79 (0.01) | 0.54 (0.02) | 0.40 (0.02) |

Since the dataset used in our work was compiled from multiple projects, it naturally encompasses a diverse range of sequences. Nonetheless, in Lead Optimization (LO) projects, new antibodies may exhibit mutations in regions that were previously unmutated in the training data, and Lead Identification (LI) projects might yield sequences that are significantly different. Assessing the robustness of the prediction pipelines in handling this challenge of out-of-distribution data is an important area for further exploration. Given the cost of measuring developability properties, our dataset of approximately 1,200 antibodies is considered moderately large. For other new or expensive low-throughput assays, we might not have access to such a large dataset, which makes it challenging to directly leverage the protein language model and graph neural network. In those cases, it would be interesting to see whether the trained models using the relatively large dataset can give an edge to learning the prediction networks for low-data assay through transfer learning technique (Golinski et al., 2023). A few recent works, e.g (Hayes et al., 2024; Malherbe & Ucar, 2024; Sun & Shen, 2024) explicitly incorporate the structural information into the training of the protein language models. Exploration of their applicability in developability prediction can provide further insights into the importance of explicit structural information in protein property prediction.

## Author Contributions

### Conceptualization

M.P., A.N.M.N.A., M.B., V.S., J.G.C., N.M., G.K., R.C.W.; **Methodology:** A.N.M.N.A., M.P.; **Data Processing:** I.S., G.K., A.N.M.N.A., M.P.; **Visualization:** A.N.M.N.A., M.P.; **Supervision:** M.P., J.S., R.C.W.; **Writing–Original Draft:** A.N.M.N.A., M.P.; **Writing– Review and Editing:** A.N.M.N.A., M.P., N.M., J.G.C., V.S., M.B.

## Acknowledgements

We would like to express our sincere gratitude to all those who have contributed to the success of this research project. Special thanks go to Jurgen Haas, Christopher Lloyd, Beverley Smith, Robert Calvert, Andrew Dippel, Bismark Amofah, and Tony Pham for providing the necessary resources and facilities to conduct this research.

## A. Cross-validation experiment for selecting best combination

### A.1. Sequence and Structure-based Features

For the prediction pipeline with sequence and structure-based features, AF2 structure prediction and Schrödinger’s protein properties have been utilized. Schrödinger provides an extensive list of protein features, not all of which may be predictive for SEC models. Consequently, a feature selection method is required to reduce the number of features. The selection process involves eliminating features with low variation (Coefficient of Variation (CV) *<* 10%), and removing highly correlated features (Spearman’s correlation *>* 90%). Additionally, we applied SHAP algorithms (Tree Explainer (Janzing et al., 2020)) using 10-fold cross-validation of the training set, repeated 100 times. The most frequently identified important features were selected for the validation dataset, and the number of features was reduced as long as the model’s performance-measured by accuracy and F1 score- remained high. These procedures reduced the number of features from an initial pool of over 1,000 to fewer than 50. Importantly, the test set was not used in any of these feature selection steps.

The majority of the selected features pertain to the hydrophobic and aggregation properties of both global and local regions of the variable fragments (Fvs). This focus makes sense because hydrophobic interactions and aggregation tendencies significantly influence the behavior of proteins during SEC (Figure 3). In SEC, proteins that aggregate or exhibit strong hydrophobic interactions often elute differently compared to those that remain monomeric and less hydrophobic, thereby affecting the resolution and accuracy of the separation process (Fekete et al., 2014; Hong et al., 2012).

**Figure 3:**
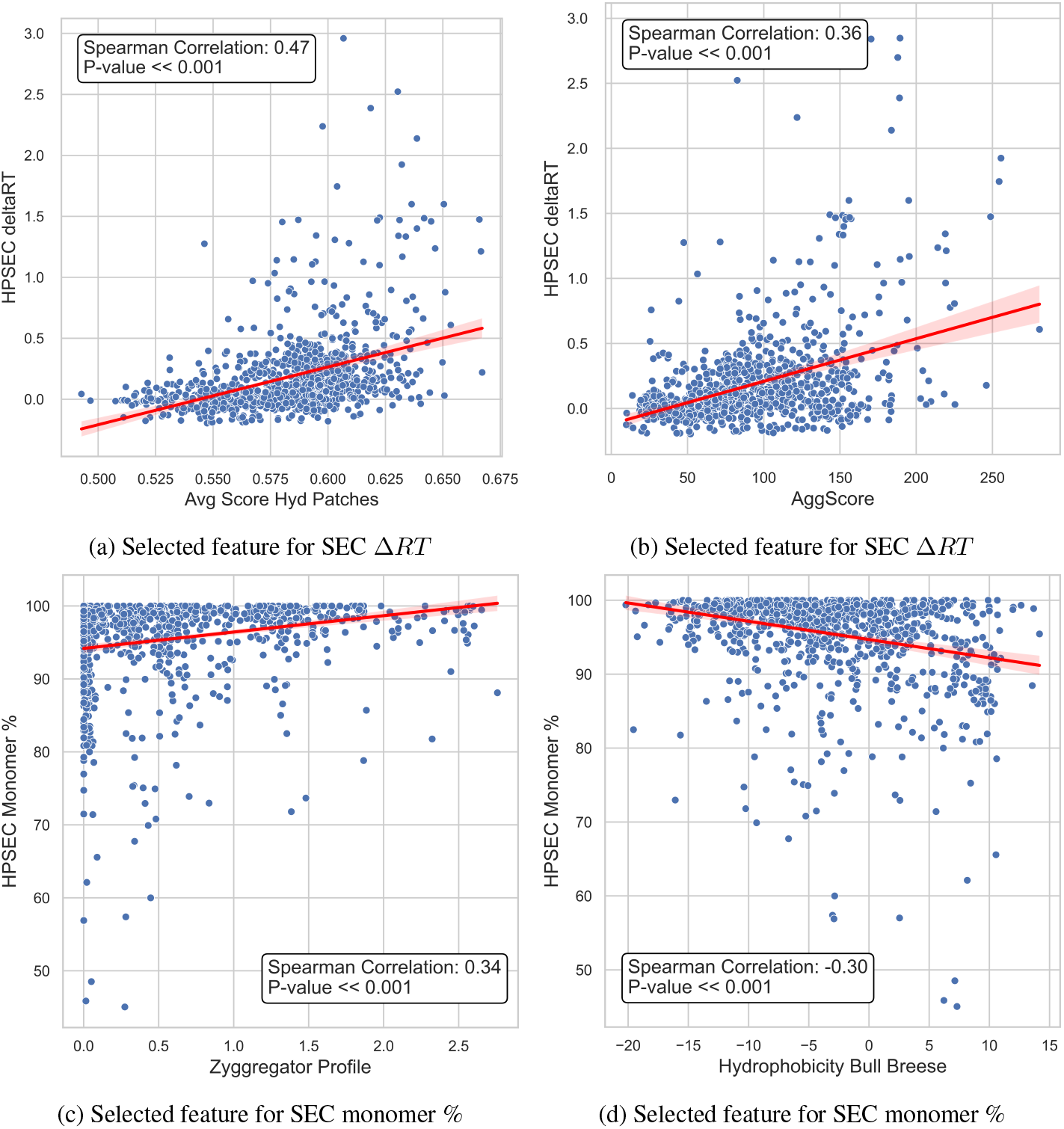
Examples of selected predictive features and their correlations to SEC attributes: The important features selected for model predictions primarily focus on the hydrophobic and aggregation properties of Fvs, as these significantly influence protein behavior and elution patterns during SEC.

### A.2. PLM, GNN and PLM+GNN pipeline

Figures 4 to 6 show the average accuracy and F1 score in the 10 trials of 10-fold cross-validation by the PLM, GNN and PLM + GNN pipelines for monomer percentage property respectively. For PLM, we have considered the pre-trained ESM2 (8M) model and learned the prediction network by either freezing its PLM weights (denoted as fixed PLM) or updating it jointly with *ϕ*_MLP_ via full parameter finetuning or LoRA. With two choices for pooling i.e. mean and CLS pooling, we have 6 combinations in total for this pipeline. In case of GNN pipeline, 3 different GNNs (GVP, GAT and GIN) with four different types of node attributes are explored. However, Figure 5 only shows the result for 9 combinations because the training with residue embedding from pre-trained ESM2 (8M) model (as node attribute) did not converge to a stable loss for monomer percentage property. Finally, 3 PLMs, 3 GNNs and 2 global pooling techniques are explored for the PLM+GNN pipeline. In each pipeline, we observed that the accuracy for different combinations is almost similar. We have selected the best-performing combination based on the F1 scores. Similar cross-validation performance for SEC Δ*RT* is shown in Figures 7 to 9.

**Figure 4:**
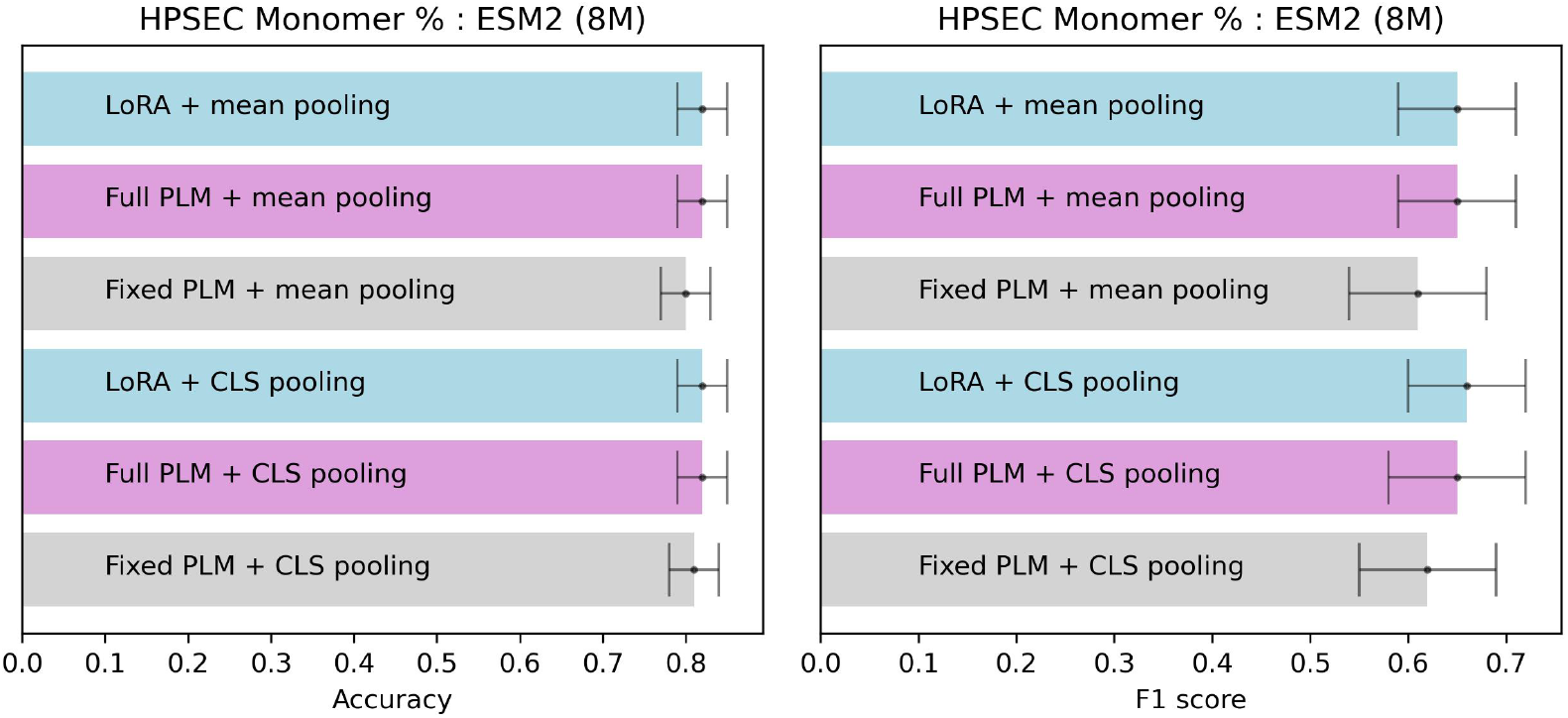
10 trials of 10 fold cross-validation performance of 6 choices of PLM pipeline for monomer %

**Figure 5:**
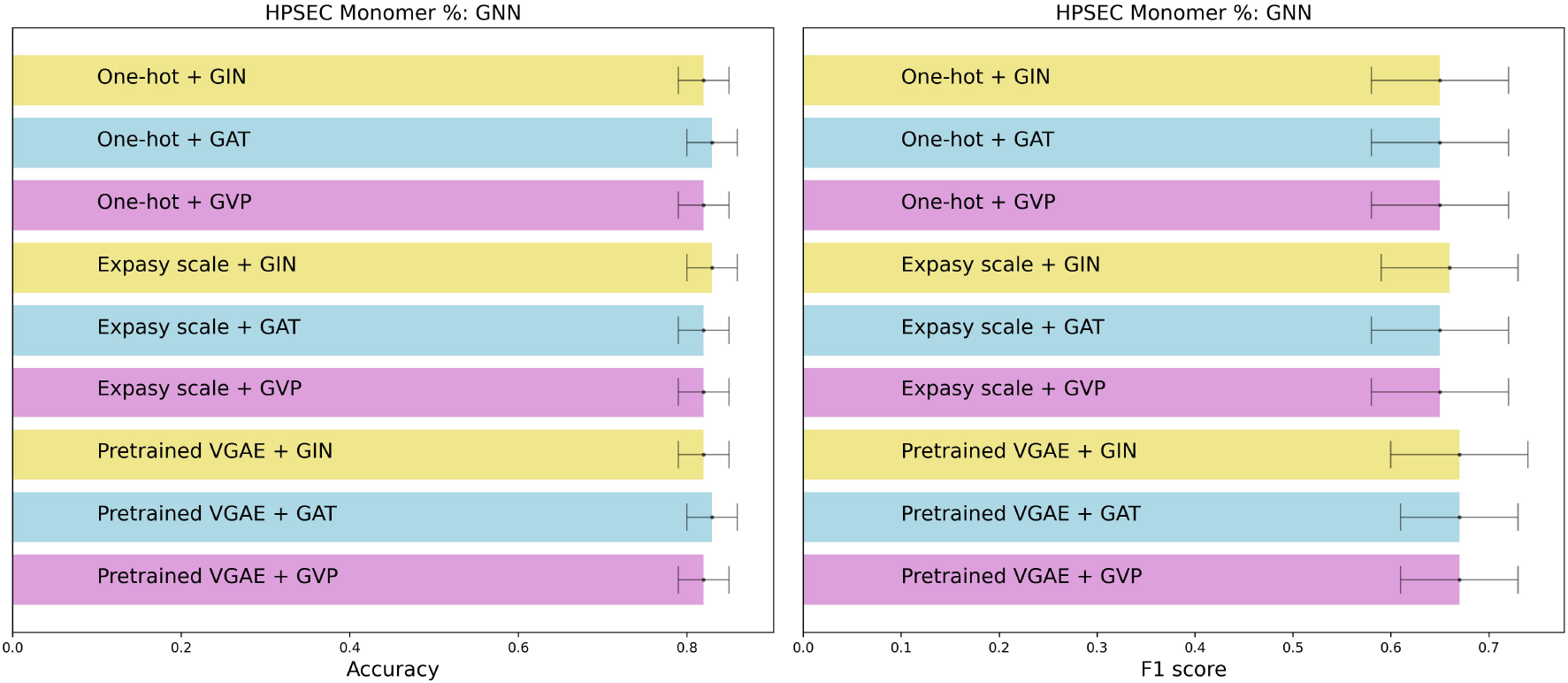
10 trials of 10 fold cross-validation performance of 9 choices of GNN pipeline for monomer %

**Figure 6:**
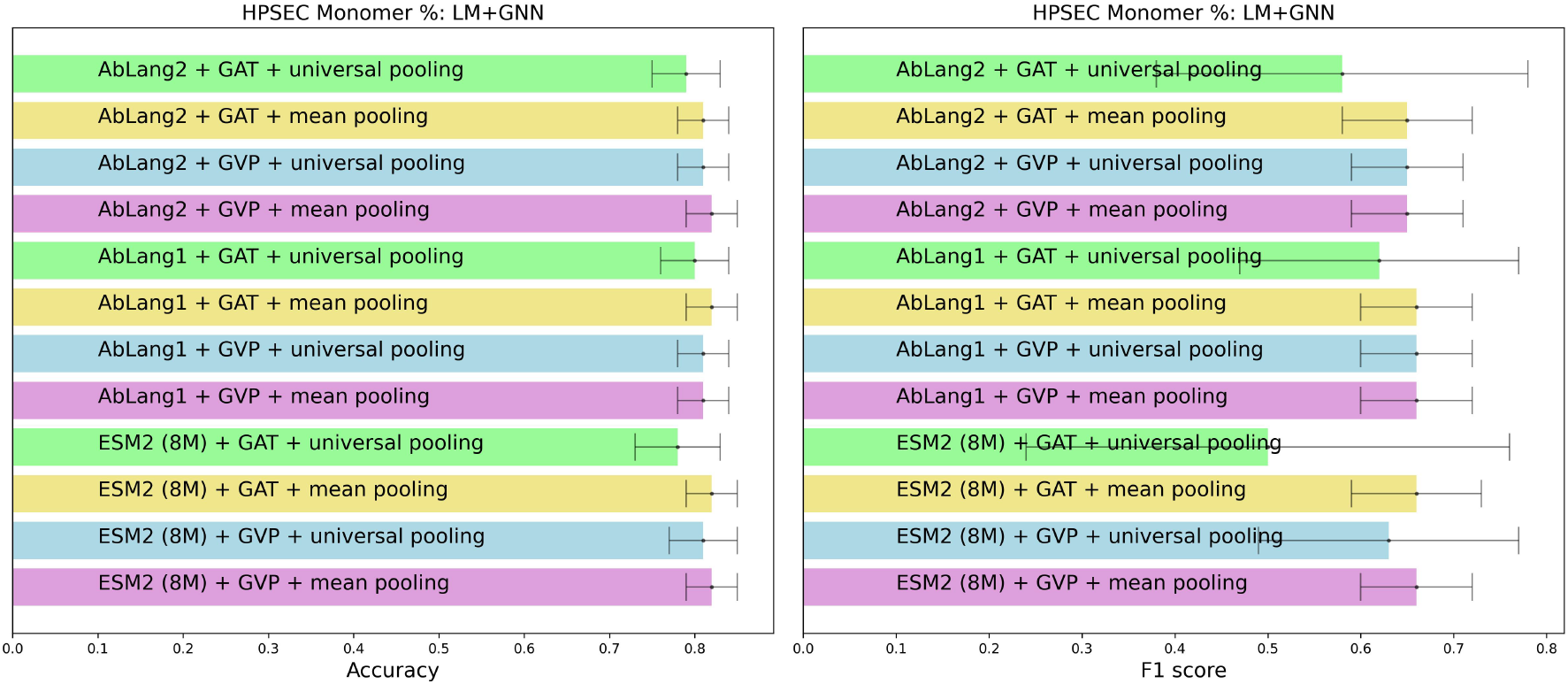
10 trials of 10 fold cross-validation performance of 12 choices of PLM+GNN pipeline for monomer %

**Figure 7:**
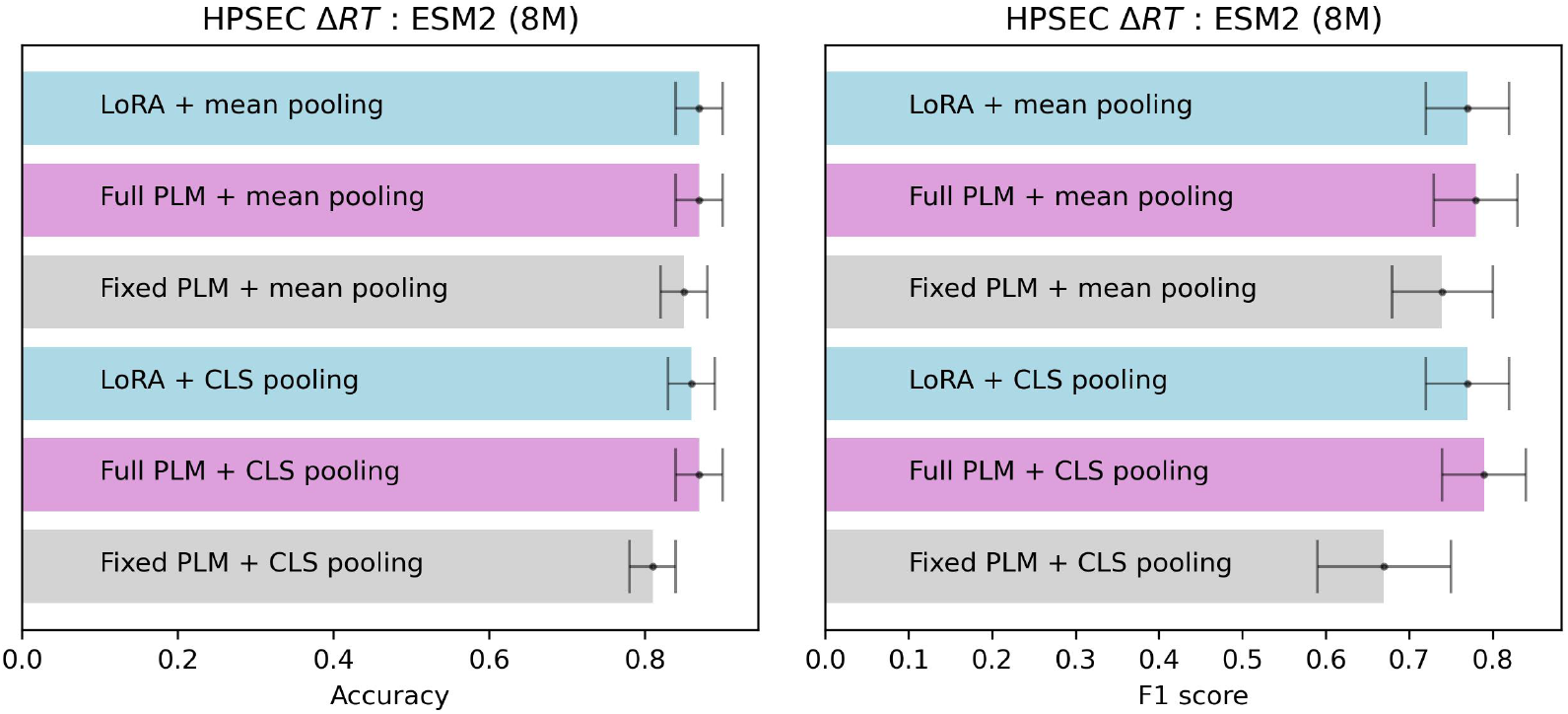
10 trials of 10 fold cross-validation performance of 6 choices of PLM pipeline for Δ*RT*

**Figure 8:**
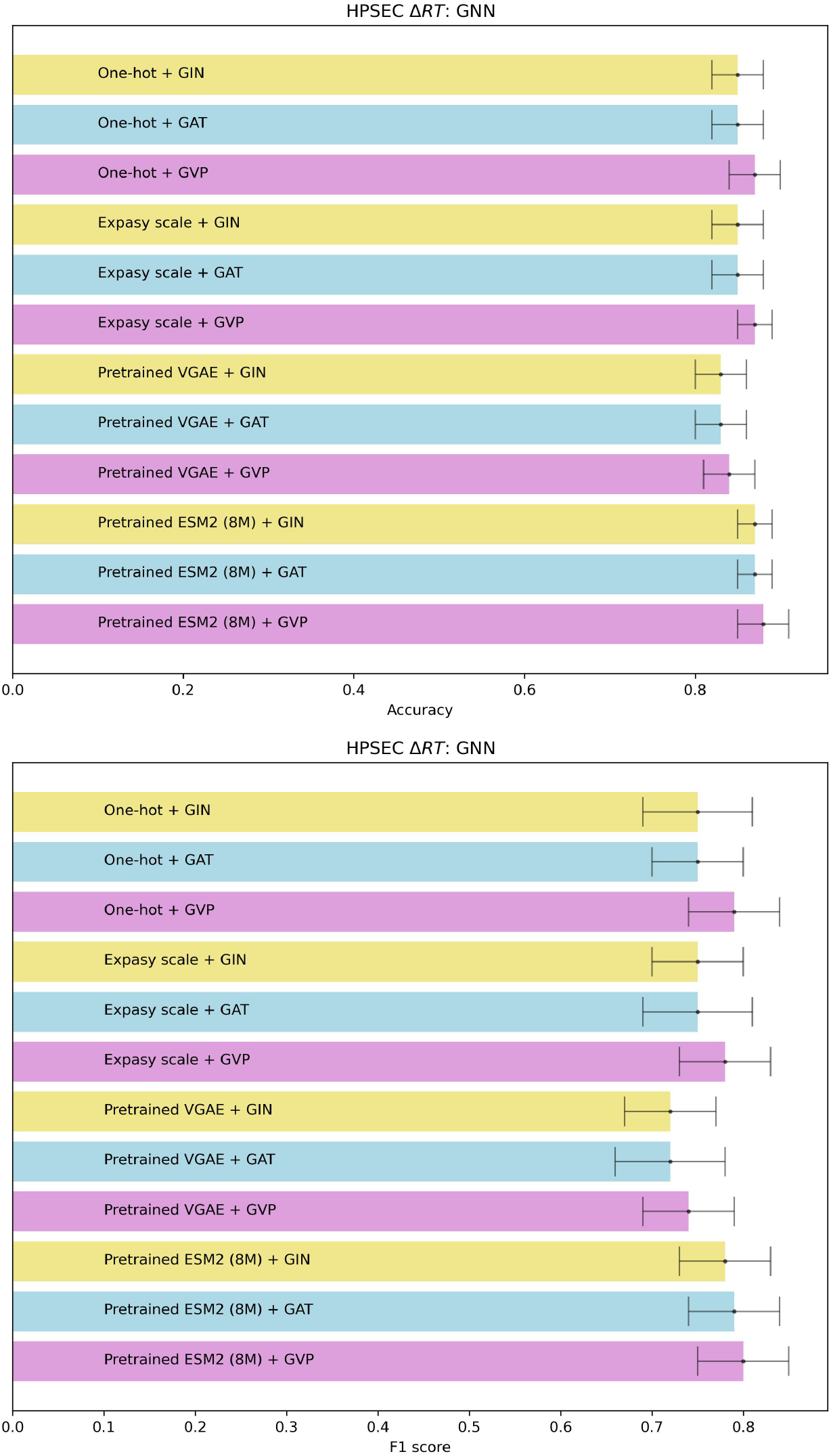
10 trials of 10 fold cross-validation performance of 12 choices of GNN pipeline for Δ*RT*

**Figure 9:**
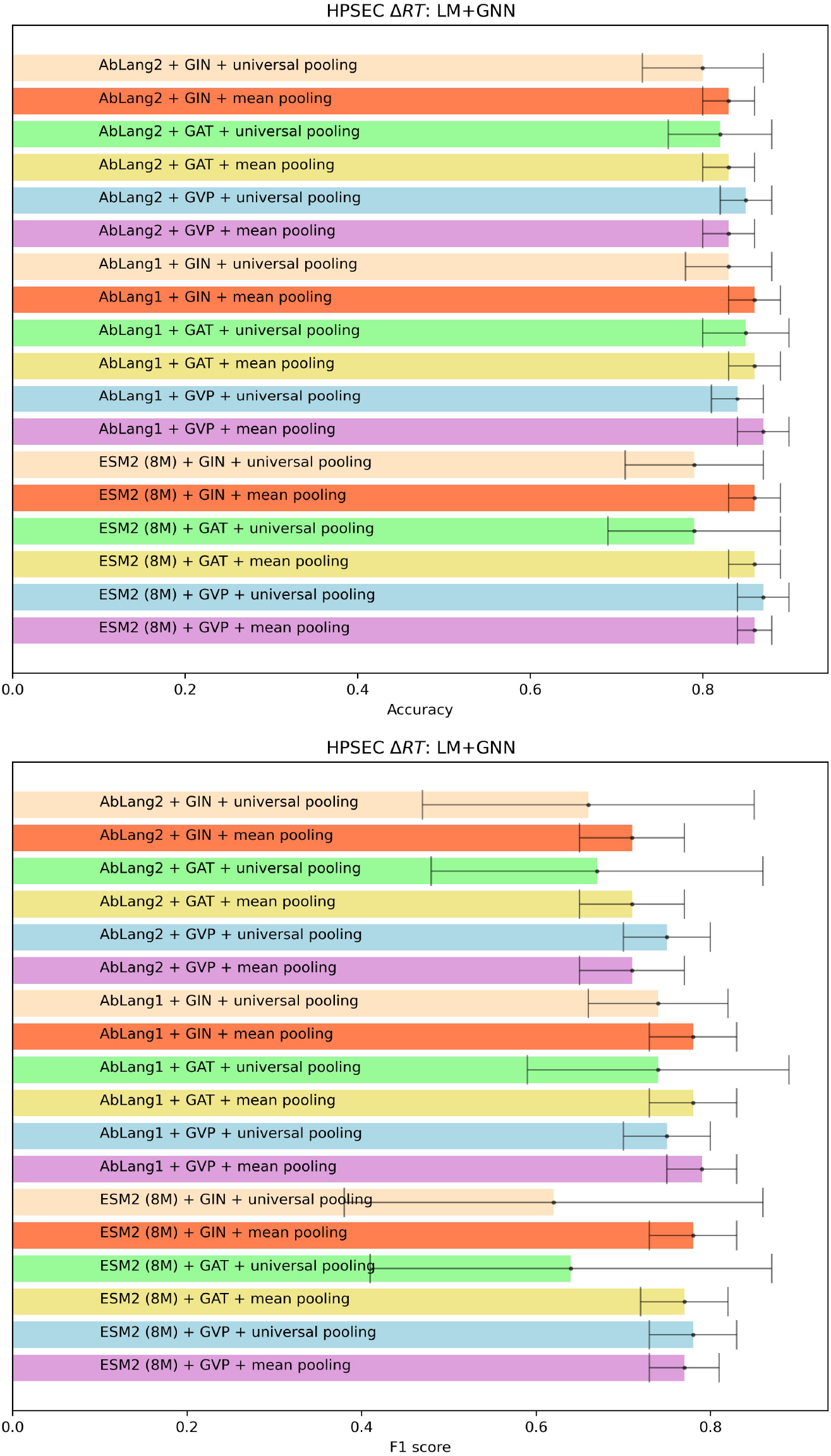
10 trials of 10 fold cross-validation performance of 18 choices of PLM+GNN pipeline for Δ*RT*

## Notes

### Competing Interest Statement

The authors have declared no competing interest.

### Summary of Updates

Author affiliations are updated on the submission portal. The manuscript is same as before.

